# Characterization of protein-ligand binding interactions of enoyl-ACP reductase (FabI) by native MS reveals allosteric effects of coenzymes and the inhibitor triclosan

**DOI:** 10.1101/2019.12.30.891283

**Authors:** P. Matthew Joyner, Denise P. Tran, Muhammad A. Zenaidee, Joseph A. Loo

## Abstract

The enzyme enoyl-ACP reductase (also called FabI in bacteria) is an essential member of the fatty acid synthase II pathway in plants and bacteria. This enzyme is the target of the antibacterial drug triclosan and has been the subject of extensive studies for the past 20 years. Despite the large number of reports describing the biochemistry of this enzyme, there have been no studies that provided direct observation of the protein and its various ligands. Here we describe the use of native MS to characterize the protein-ligand interactions of FabI with its coenzymes NAD^+^ and NADH and with the inhibitor triclosan. Measurements of the gas-phase affinities of the enzyme for these ligands yielded values that are in close agreement with solution-phase affinity measurements. Additionally, FabI is a homotetramer and we were able to measure the affinity of each subunit for each coenzyme, which revealed that both coenzymes exhibit a positive homotropic allosteric effect. An allosteric effect was also observed in association with the inhibitor triclosan. These observations provide new insights into this well-studied enzyme and suggest that there may still be gaps in the existing mechanistic models that explain FabI inhibition.

## Introduction

Around 20 years ago a flurry of reports appeared in the scientific literature, all describing various structural and functional details of the bacterial enzyme enoyl-ACP reductase (FabI) from *E. coli*.^1–5^ The excitement surrounding this enzyme had been ignited by the discovery that it was the biological target for two widely used antibacterial agents, triclosan and isoniazid.^6–8^ Triclosan has been used extensively as an active ingredient in hand soaps and other personal care products for the past 50 years, but it was believed to act directly on bacterial membranes and therefore have no specific biomolecular interactions.^7,8^ Isoniazid has been used as a treatment for tuberculosis for many decades,^6,9^ and the discovery that both of these compounds inhibited the same enzyme led many in the scientific community to begin investigating enoyl-ACP reductase as a candidate target for new antibacterial therapeutics.

Although much of the interest in enoyl-ACP reductase as a drug target has cooled, extensive work has been published describing its biochemical mechanism and structural features. ^5,10–13^ The enzyme catalyzes the final reductive step in the bacterial fatty acid synthesis cycle, with NAD(P)H providing a hydride that adds to C3 of the enoyl substrate followed by the addition of a proton from water to the enolate intermediate (Figure 1).

**Figure 1.**
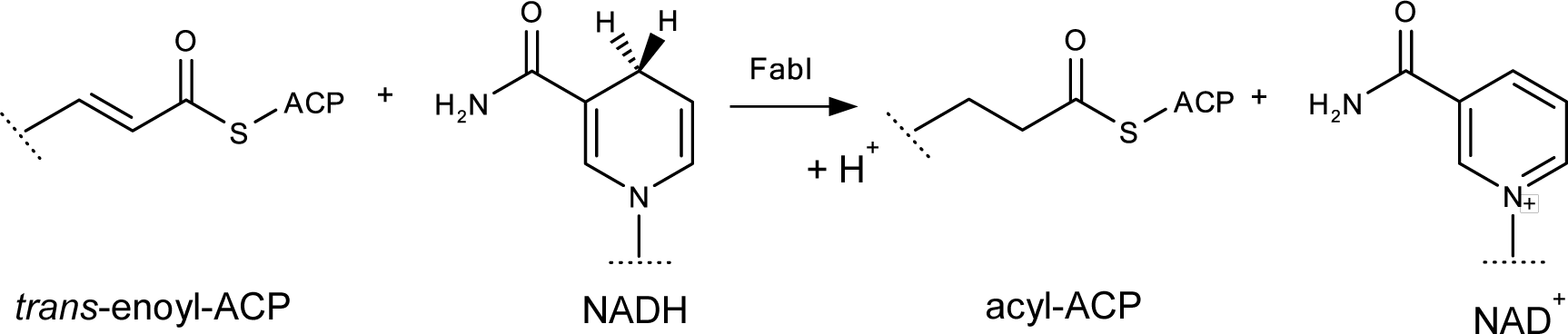
The enzyme enoyl-ACP reductase in *E. coli* (FabI) catalyzes the reduction of an enoyl group covalently linked to an acyl carrier protein (ACP), with NAD(P)H acting as a coenzyme hydride donor.

The product dissociates from the active site first, followed by dissociation of the oxidized coenzyme (NAD^+^).^14,15^ The drug triclosan has been reported to act as a slow-binding inhibitor of the enzyme by reversibly binding in the active site on top of the NAD^+^ and forming an inactive ternary complex (Figure 2).

**Figure 2.**
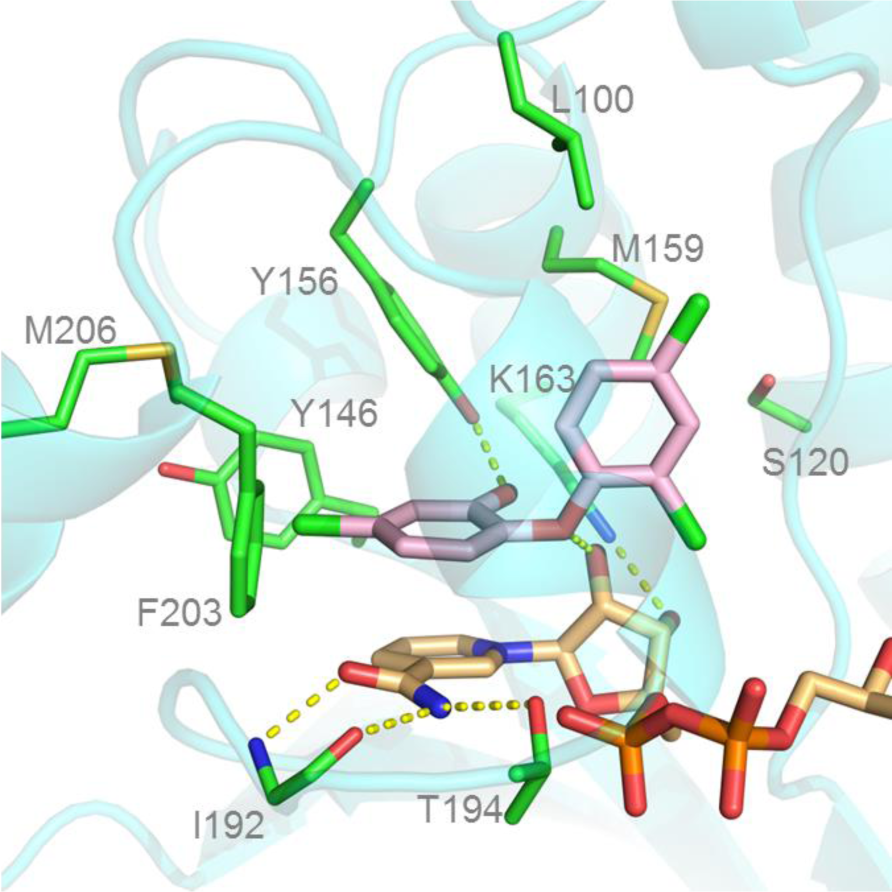
The enoyl-ACP reductase active site (PDB 1QSG). Residues that form the active site and participate in catalysis are labeled. Hydrogen bonding interactions in the active are shown as yellow dashed lines: the 2ʹ-OH of triclosan (magenta) with Y156 and the bridging oxygen with the 2ʹ-OH of the ribosyl of the ribosyl of NADH; the amide group of the nictotinamide ring in NAD^+^ (orange) with main chain atoms of I192 and the O^γ^ of T194, and the 3ʹ-OH of the ribosyl with the N^ε^ of K163.

This mechanism has been supported by many x-ray crystal structures that show triclosan in the active site, forming H-bonds with the catalytic Y156 residue and stabilizing a flexible loop in a closed conformation over the triclosan binding site.^1–5^

In this report, we have used native protein nanosprayESI mass spectrometry to characterize the physical interactions of FabI with the ligands NADH, NAD^+^, and triclosan. Native MS has been repeatedly demonstrated to provide meaningful and unique information about protein structures and is particularly useful for the interrogation of protein-ligand binding interactions.^16–20^ Native MS has also been demonstrated as a unique tool for the investigation of allosteric properties of proteins since the protein-ligand interactions can be directly observed rather than inferred from indirect measurements.^21^ Our measurement of the binding affinity of NADH and NAD^+^ with FabI are in line with previous reports from solution-phase studies. However, our data suggests that triclosan induces a stable change in the tertiary structure of FabI that results in a very large increase in the enzyme’s affinity for NAD^+^, and that this structural change may be stable even after triclosan dissociates from the enzyme.

## Results

FabI was overexpressed from a plasmid in *E. coli* and isolated using Ni-affinity chromatography. Native MS analysis of 5 µM solutions of the purified enzyme clearly showed the FabI apo tetramer at charge states of +22, +21, and +20, with the +21 charge state being the most abundant (Figure 3A). In addition to the apo enzyme, a small amount of FabI-ligand complex was observed, despite repeated washings through both molecular weight cutoff filters and size-exclusion spin columns. Deconvolution of the apo spectrum yielded a measured mass of 112,613.62 ± 31.72 Da for the apo peak, which was very close to the expected mass of 112,580 Da. The deconvoluted mass of the small peak was 113,300.10 ± 26.49 Da, which is consistent with the mass of a FabI-NAD^+^ complex; the identity of the complex was confirmed by the observation of peaks with the expected mass of NAD^+^ that were observed in a denatured sample of FabI (Figure S1). Addition of 64 µM NADH to the FabI resulted in the observation of FabI-NADH complexes with 1, 2, 3 and 4 ligands bound to the enzyme (Figure 3B). At higher concentrations of NADH, complexes with 5 and 6 ligands bound were also observed.

**Figure 3.**
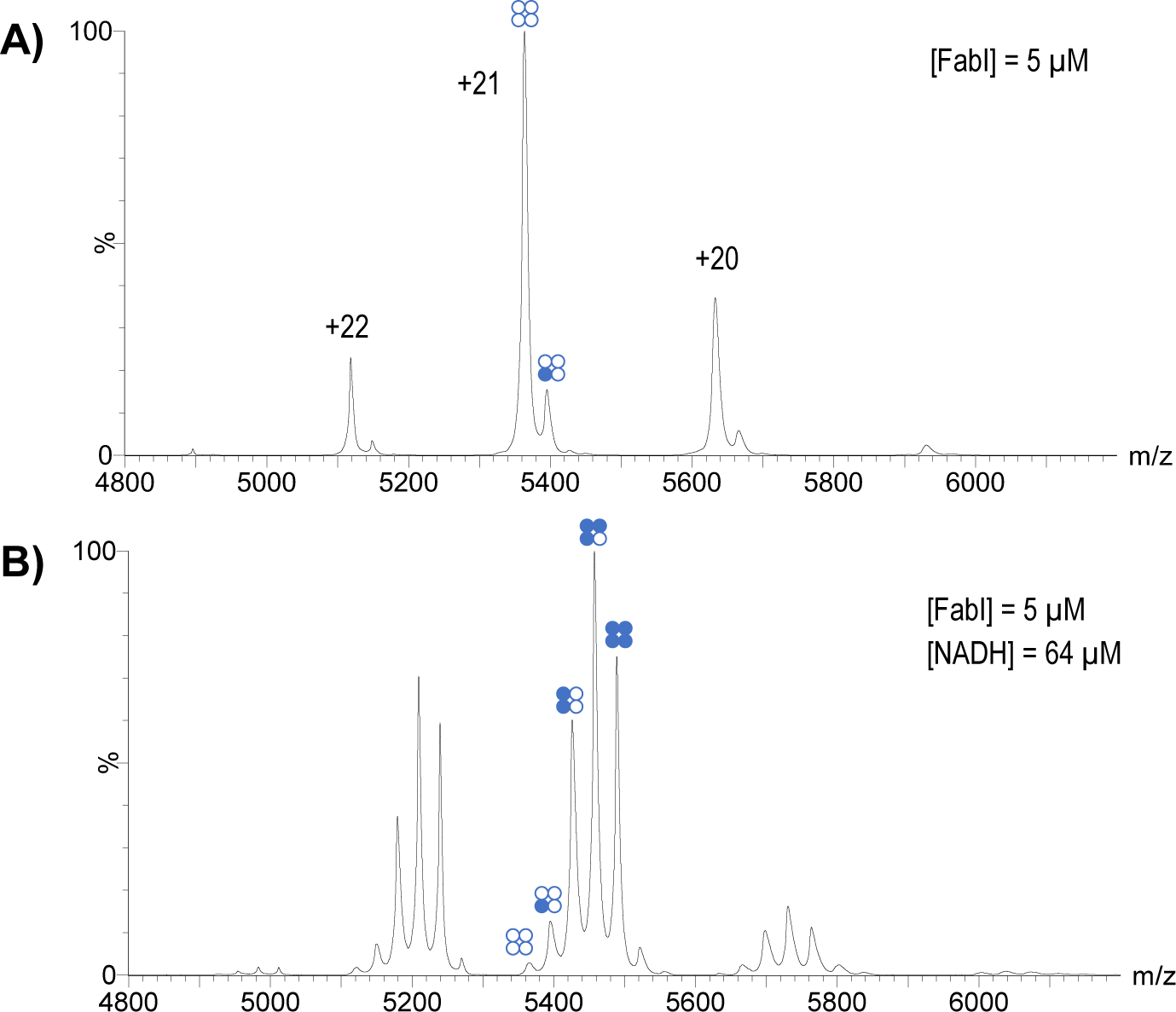
Native mass spectra of FabI tetramer. Charge states are shown and bubble diagrams indicate apo (empty circles) and holo forms (filled circles) of the enzyme. A) FabI (5 µM) with no ligand added. Small amount of holo-enzyme could not be separated from apo enzyme. B) FabI (5 µM) with NADH (64 µM). The apo-enzyme has almost completely disappeared and holo-enzyme with 1, 2, 3, and 4 NADH ligands bound are observed.

Treatment of FabI with 64 µM NAD^+^ caused an increase in the intensity of the peaks corresponding to the FabI-NAD^+^ holo complex with a single ligand bound and the appearance of very small amounts of a complex with 2 ligands bound (Figure 4A). Addition of triclosan to FabI resulted in a negative shift in the charge state distribution, with the +20 charge state being the most abundant (Figure 4B). Although no NAD^+^ was added to this solution, the intensity of the peak corresponding to the FabI-NAD^+^ complex increased, relative to the intensity of the FabI apo peak. No peaks corresponding to a FabI-triclosan complex were observed. We hypothesized that the absence of this observation was caused by in-source dissociation of triclosan from the enzyme, but we were still unable to observe the FabI-triclosan complex even after extensive effort to optimize the ionization parameters.

**Figure 4.**
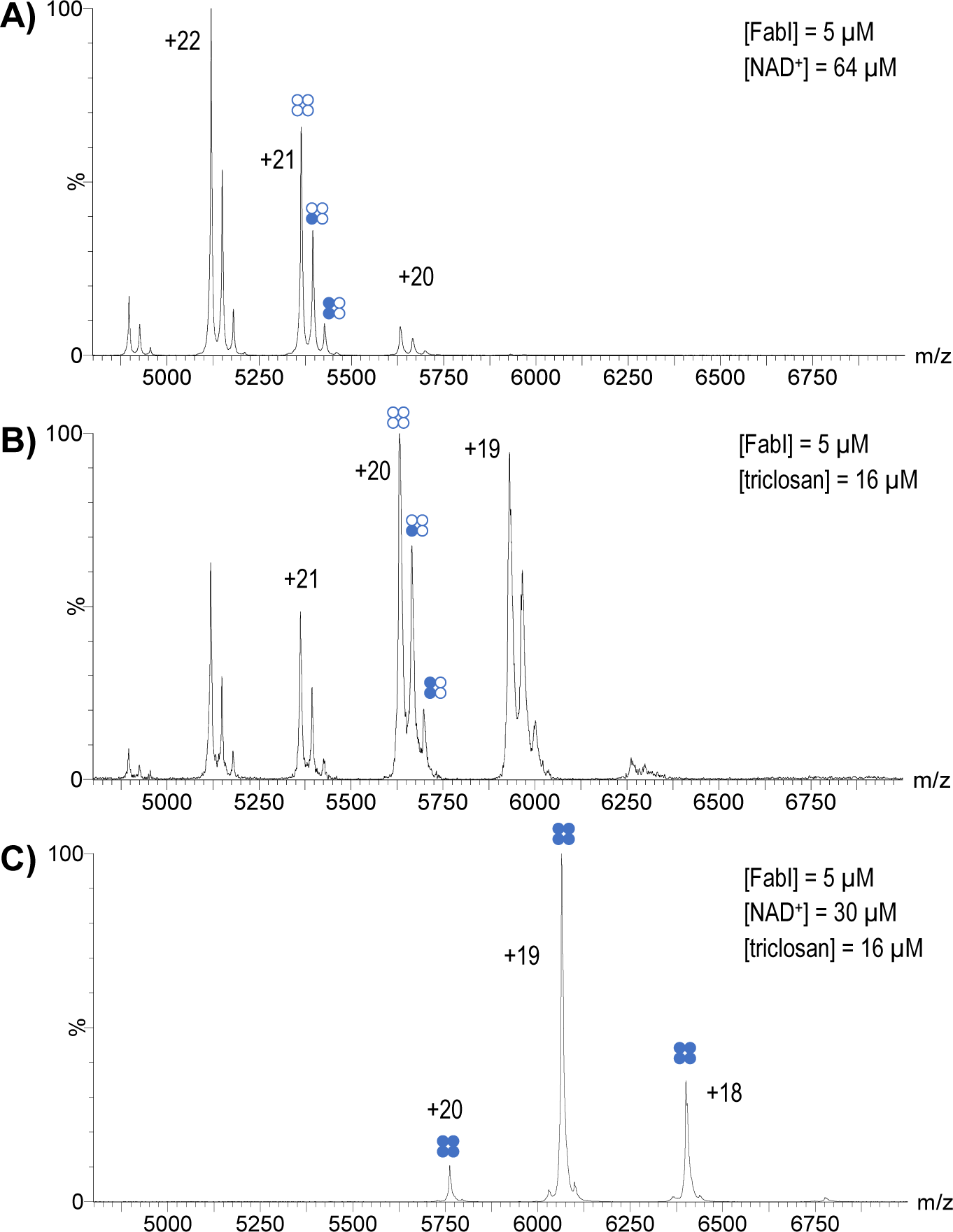
Native MS of A) Fabi-NAD^+^, B) FabI-triclsoan, and C) FabI-NAD^+^-triclosan. A) FabI (5 µM) with NAD^+^ (40 µM). The majority of the enzyme is in the apo form, but holo enzyme with 1 and 2 NAD^+^ ligands bound can be observed. B) FabI (5 µM) with triclosan (8 µM). Addition of triclosan caused a negative shift in the charge state distribution and a visible increase in the relative amount of holo enzyme. No NAD^+^ was added to this sample, so presumably the increased intensity of the holo enzyme arises from the endogenous protein-ligand complex that could not be removed from the sample. C) FabI (5 µM) with NAD^+^ (30 µM) and triclosan (16 µM). Only the holo enzyme with 4 NAD^+^ ligands bound can be observed.

Solution-phase and x-ray crystallography studies of FabI have suggested that triclosan inhibits FabI by blocking NAD^+^ release from the active site after completion of a catalytic cycle.^1,5^ Therefore, we prepared a mixture of FabI (5 µM), NAD^+^ (30 µM) and triclosan (16 µM) and observed only a single peak, corresponding to a FabI-4NAD^+^ complex (Figure 4C). This observation presumably is caused by a dramatic increase in the binding affinity of FabI for NAD^+^. The complete disappearance of peaks corresponding to the apo enzyme indicates that all of the subunits of each enzyme are functional and capable of binding the coenzyme.

Titrations of NAD^+^ and NADH with FabI were performed to measure the binding equilibrium association constants (K_A_) for each ligand. Using equations derived by John Klassen and colleagues,^22^ we calculated the macroscopic K_A_ for each ligand and microscopic K_A_ values for each subunit in the tetramer (Table 1).

**Table 1.**
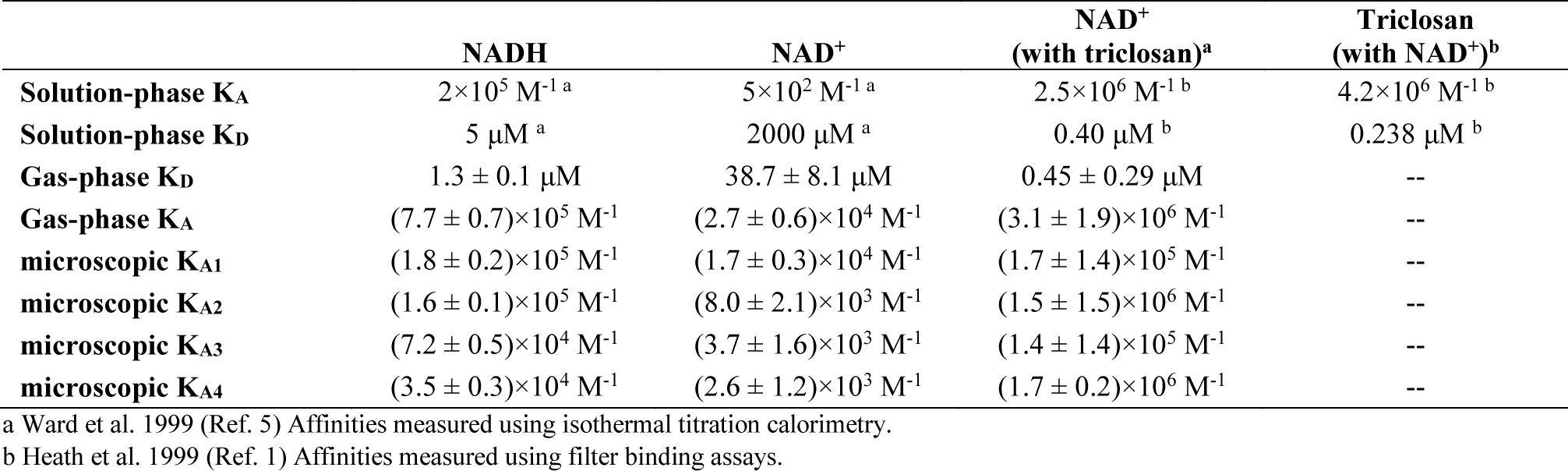
Summary of FabI binding affinity values for NADH and NAD^+^ and triclosan.

|  | NADH | NAD<sup>+</sup> | NAD<sup>+</sup><br>(with triclosan)<sup>a</sup> | Triclosan<br>(with NAD<sup>+</sup>)<sup>b</sup> |
| --- | --- | --- | --- | --- |
| **Solution-phase K<sub>A</sub>** | 2×10<sup>5</sup> M<sup>-1</sup> <sup>a</sup> | 5×10<sup>2</sup> M<sup>-1</sup> <sup>a</sup> | 2.5×10<sup>6</sup> M<sup>-1</sup> <sup>b</sup> | 4.2×10<sup>6</sup> M<sup>-1</sup> <sup>b</sup> |
| **Solution-phase K<sub>D</sub>** | 5 μM <sup>a</sup> | 2000 μM <sup>a</sup> | 0.40 μM <sup>b</sup> | 0.238 μM <sup>b</sup> |
| **Gas-phase K<sub>D</sub>** | 1.3 ± 0.1 μM | 38.7 ± 8.1 μM | 0.45 ± 0.29 μM | -- |
| **Gas-phase K<sub>A</sub>** | (7.7 ± 0.7)×10<sup>5</sup> M<sup>-1</sup> | (2.7 ± 0.6)×10<sup>4</sup> M<sup>-1</sup> | (3.1 ± 1.9)×10<sup>6</sup> M<sup>-1</sup> | -- |
| **microscopic K<sub>A1</sub>** | (1.8 ± 0.2)×10<sup>5</sup> M<sup>-1</sup> | (1.7 ± 0.3)×10<sup>4</sup> M<sup>-1</sup> | (1.7 ± 1.4)×10<sup>5</sup> M<sup>-1</sup> | -- |
| **microscopic K<sub>A2</sub>** | (1.6 ± 0.1)×10<sup>5</sup> M<sup>-1</sup> | (8.0 ± 2.1)×10<sup>3</sup> M<sup>-1</sup> | (1.5 ± 1.5)×10<sup>6</sup> M<sup>-1</sup> | -- |
| **microscopic K<sub>A3</sub>** | (7.2 ± 0.5)×10<sup>4</sup> M<sup>-1</sup> | (3.7 ± 1.6)×10<sup>3</sup> M<sup>-1</sup> | (1.4 ± 1.4)×10<sup>5</sup> M<sup>-1</sup> | -- |
| **microscopic K<sub>A4</sub>** | (3.5 ± 0.3)×10<sup>4</sup> M<sup>-1</sup> | (2.6 ± 1.2)×10<sup>3</sup> M<sup>-1</sup> | (1.7 ± 0.2)×10<sup>6</sup> M<sup>-1</sup> | -- |

<sup>a</sup> Ward et al. 1999 (Ref. 5) Affinities measured using isothermal titration calorimetry.
<sup>b</sup> Heath et al. 1999 (Ref. 1) Affinities measured using filter binding assays.

Consistent with previous solution-phase studies of FabI, measurements of the macroscopic affinities accounting for the combination of all binding sites showed that NAD^+^ exhibited an approximately 28-fold weaker affinity for FabI than NADH, with values of (2.7 ± 0.6)×10^4^ M^−1^ and (7.7 ± 0.7)×10^5^ M^−1^, respectively. Measurements of the microscopic affinity of each individual binding site for the two ligands showed a progressing decrease in affinity for subsequent binding events. Based on the formula for calculating the statistical relationship between the microscopic and macroscopic K_A_ values^22^ (equation 4), the individual binding affinities are not independent, suggesting cooperativity among the subunits. The observed microscopic K_A_ values for NADH ranged from 16-fold to 4-fold lower than expected for independent binding sites, while the microscopic K_A_ values for NAD^+^ ranged from 5-fold to 1.5-fold lower than expected for independent sites. For both ligands, the large deviation from the expected numerical relationship between the macroscopic and microscopic binding affinities provided indicates that both ligands exhibit a positive homotropic allosteric effect.

Treatment of the enzyme with triclosan caused a ~100-fold increase in affinity of FabI for NAD^+^ (Table 1), which is generally consistent with affinity measurements from solution-phase membrane-binding assays reported by Charles Rock and coworkers.^1^ The reported solution-phase affinity for NAD^+^ is ~50-fold higher than we measured in the gas-phase in this study; the solution-phase studies reported difficulty in obtaining accurate measurements for this affinity value^1,5^ which makes it difficult to compare our measured change in affinity for NAD^+^ to these prior studies. We also measured the microscopic NAD^+^ K_A_ values in the presence of triclosan for each binding site, which revealed that the second and fourth binding sites have approximately 10-fold greater affinity for NAD^+^ than the first and third binding sites (Table 1), suggesting a possible cooperative interaction between each symmetric dimer in the homotetramer. Additionally, comparison of the measured values to the expected values showed that K_1_ is ~70-fold less than expected and K_4_ is ~2-fold greater than expected, which is consistent with our observed shift of FabI to being completely occupied by NAD^+^ in the presence of triclosan (Figure 4B).

To evaluate the relative stability of the changes to FabI induced by triclosan we used CID to dissociate the tetramer to the monomer. Application of 90V at the ion trap caused a large degree of dissociation and clearly visible monomer peaks with charge states ranging from +15 to +10 (Figure 5). MS/MS experiments confirmed that the higher charge state monomers were primarily dissociation products of the higher charge state tetramers and lower charge state monomers were primarily dissociation products of lower charge state tetramers (Figure S2). With the addition of 30 µM NAD^+^, only the apo monomer was visible with no peaks corresponding to holo complexes observed (Figure 5A). However, when 16 µM triclosan was added to the solution, in addition to the apo monomer a holo monomer was also observed even with application of 90V CID energy (Figure 5B). The mean mass difference of the holo monomeric complex compared to the apo monomer was 540.892 Da, which is a good match for the calculated mass of a NAD^+^ fragment product of the cleavage of the nicotinamide-ribosyl glycosidic bond (540.052 Da). MSMS experiments confirmed that the monomeric FabI-ligand complex was a dissociation product from the tetramer (Figure S3). The monomeric peaks corresponding to the FabI-NAD^+^ fragment complex is also shifted to a lower charge state distribution, similar to the shift observed in the native spectra of the tetramer. Thus, the changes in FabI induced by triclosan were stable in the gas phase even after dissociation of the tetramer.

**Figure 5.**
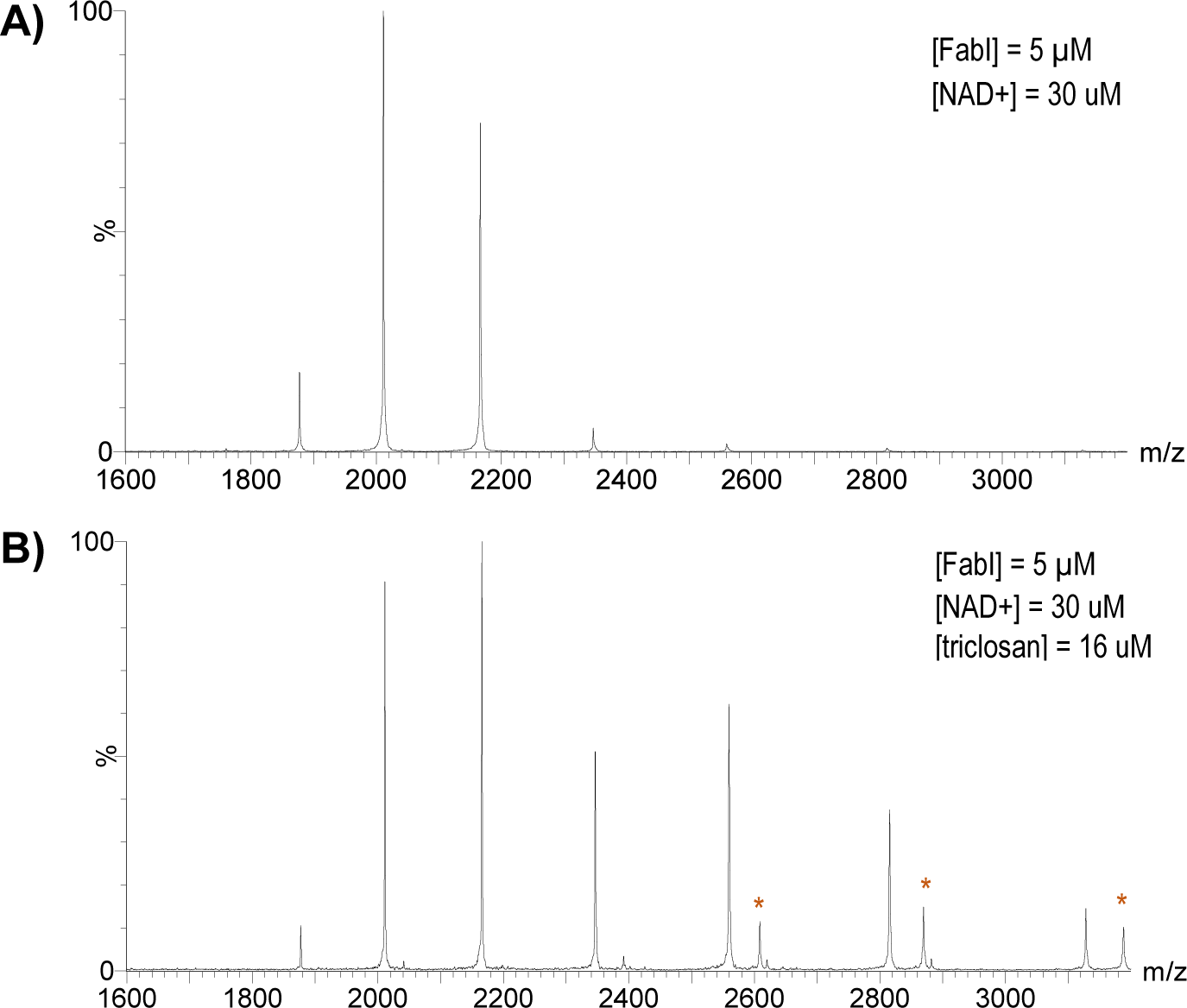
CID (90V) MS spectra of A) Fabi-NAD^+^, and B) FabI-NAD^+^-triclosan. A) FabI (5 µM) with NAD^+^ (30µM). Only the apo monomer can be observed. B) FabI (5 µM) with NAD^+^ (30 µM) and triclosan (16 µM). A distribution of monomers that matches the FabI-NAD^+^ spectra is present, but a second distribution with lower charge states is also observed. A peak corresponding to a FabI-ligand monomeric complex is also visible in the distribution with lower charge states (marked with *).

The observation of the monomeric FabI-ligand complex suggested that treatment with triclosan induced a conformational change that led to a strong FabI-NAD^+^ binding interaction even in the absence of triclosan. To assess this hypothesis, a single-time point hydrogen-deuterium exchange experiment was performed. For each sample, FabI, NAD^+^, triclosan and 500 mM ammonium acetate were diluted into D_2_O (D_2_O >90% for all samples). After equilibrating the mixtures for 10 minutes, MS spectra were collected (Figure 6). Samples equilibrated in D_2_O (Figure 6B, D) showed an obvious shift to higher mass for each charge state, compared to samples collected in H_2_O (Figure 6A, C). The degree of exchange increased as charge state decreased, so the degree of exchange was compared for samples with and without triclosan at only the +20 charge state. The mass shift was calculated to be 43.374 m/z units without triclosan and 36.8975 m/z units with the triclosan treatment.

**Figure 6.**
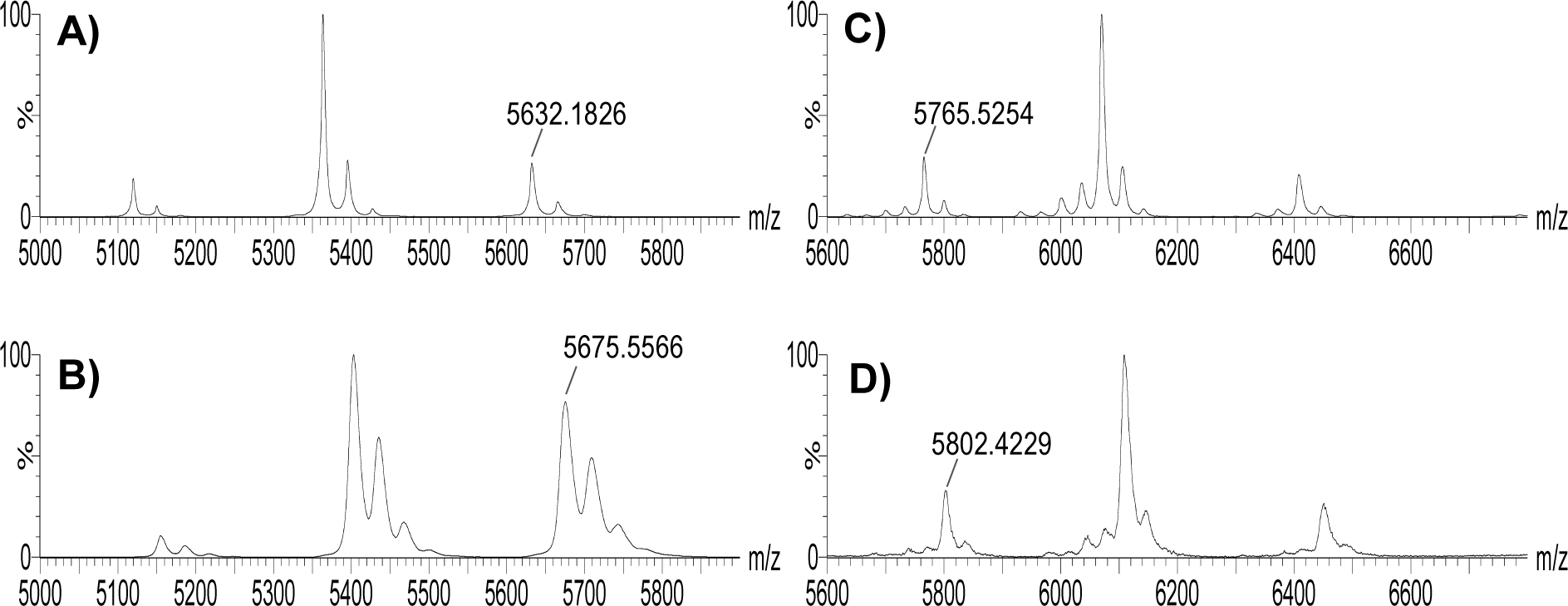
Single-time point hydrogen exchange experiment with panels A) and C) showing the spectra collected in H_2_O and panels C) and D) showing the spectra collected in D_2_O. The m/z value of the +20 charge state is shown in each panel. A) FabI (5 µM) with NAD^+^ (30 µM) in H_2_O. B) FabI (5 µM) with NAD^+^ (30 µM) in D_2_O. C) FabI (5 µM) with NAD^+^ (30 µM) and triclosan (16 µM) in H_2_O. D FabI (5 µM) with NAD^+^ (30 µM) and triclosan (16 µM) in D_2_O.

## Discussion

Although MS yields observations of biomolecules in the gas phase, the use of native protein MS to investigate protein-ligand interactions has been repeatedly demonstrated to generate information that is applicable to solution-phase chemistry of proteins.^18–21^ The great advantage provided by native MS studies is the ability to directly observe protein-ligand interactions rather than inferring them from indirect measurements of kinetics or other properties. Our observations of FabI in this report confirm much of what has previously been published about this enzyme, and yet reveals new insights that may open new paths to the development of inhibitors of this and related enzymes. First, our measurements of FabI and its affinity for the coenzyme NADH confirm that it binds to the enzyme with a 1:4 protein:ligand stoichiometry, consistent with its homotetrameric structure (Figure 3). Our binding affinity measurements are also in general agreement with the relative affinities measured in solution-phase experiments (Table 1). Our measurements show an approximately 28-fold difference in affinity, while solution-phase experiments reported a 40-fold difference in affinity.^5^ These results can be considered a rough match, especially because in the 1999 study by Ward et al., the authors state that they had difficulty generating accurate results for their affinity measurements.

The microscopic K_A_ values we measured demonstrated positive cooperativity in binding of both NADH and NAD^+^ since the statistical relationship between the microscopic and macroscopic K_A_ values failed to follow the expected pattern and since homotropic effects must be positive by definition.^23^ Positive cooperativity of NADPH for FabI from *S. aureus* has been previously observed,^24,25^ but this is the first report of cooperativity for *E. coli* FabI. It is surprising that this cooperativity has not been reported previously, but it is likely due to the difficulty of measuring NADH/NAD^+^ binding affinities using solution-phase methods that have been described by the authors of prior studies.^5,24^

The observation of the FabI-NAD^+^ complex in purified samples was surprising considering that solution-phase studies of FabI have reported relatively low affinity for NAD^+^ (Table 1);^1,5^ it is unclear if this observation has any functional significance. A dramatic change in the MS spectrum of FabI was observed when triclosan was added to the enzyme with NAD^+^ (Figure 4C). Under these conditions, almost all of the enzyme shifted to a holo form with 4 NAD^+^ ligands bound. This result demonstrates that all of the active sites in all of the tetramer molecules in solution are capable of binding NAD^+^ and that treatment with triclosan induces a change in FabI that pushes the equilibrium toward the completely occupied state. Hypotheses based on x-ray crystal structures suggest that triclosan binding in the active site induces a conformational change in FabI that stabilizes a flexible loop in a closed conformation over the FabI-NAD^+^-triclosan ternary complex.^1–5^ Our MS experiments do not present any evidence that would negate these hypotheses, however our measured microscopic K_A_ values for NAD+ in the presence of triclosan (Table 1) show a clear departure from the expected values for independent binding events. The pattern of changes in affinity in these miscroscopic K_A_ values are also very different from the patterns observed for the coenzymes alone. Without triclosan, each coenzyme exhibits a homotropic effect with progressively decreasing affinities (Table 1). In the presence of triclosan, the values for K_2_ and K_4_ are ~10-fold greater than the values for K_3_ and K_4_, which suggests that triclosan causes a conformational change that has unequal effects on the four binding sites. Therefore, it appears that triclosan is also exerting an allosteric effect on FabI as part of its inhibitory mechanism. This is a new observation for this enzyme and will require further experiments to understand its full implications.

The HX experiments (Figure 6) showed that there is a conformational change induced by triclosan, since decreased exchange was observed with triclosan treatment. A decrease in hydrogen exchange would suggest an increase in the protection of amide protons through the formation of stable hydrogen-bonding partners ^26^. The formation of more defined secondary structure because of triclosan binding would be consistent with X-ray crystal structures that show a significant unstructured region near the active site that becomes an ordered α-helix in the presence of triclosan.^3^ Additionally, examination of the FabI-triclsoan-NAD ternary x-ray crystal structure (PDB 1QSG) shows that when the “flipping loop” of FabI closes over the triclosan binding pocket, K201 is positioned so that it can form a salt bridge with D101. Since salt bridges are expected to exhibit a greater stabilizing effect in the gas phase,^27^ the formation of this salt bridge could explain why the effect of triclosan on FabI is still observed in our MS experiments even after triclosan has been ejected from the enzyme. However, some conformational freedom appears to remain, as suggested by our observation of the NAD^+^ fragment that remains bound to the monomer even after it has dissociation from the tetramer (Figure 5). The formation of a salt bridge between a FabI basic residue and the phosphoester group of NAD^+^ could explain this strong interaction, but examination of the X-ray structure does not place any basic side chains closer than 11 Å. The best candidate residue to participate in this type of conformational rearrangement would probably be R193, since it is located on the flexible loop that is stabilized by triclosan. A rotation of the peptide backbone of the flexible loop could orient the arginine towards NAD^+^, thus providing a mechanism for salt bridge formation. On the surface this appears implausible, but there are no other apparent residues in the flexible loop that could form strong enough interactions with NAD^+^ to explain our observations. An alternative explanation could be proposed that makes use of a conformational change in a different part of FabI away from the flexible loop. However, no evidence exists in a multitude of x-ray structures for extensive conformational flexibility in other parts of the FabI structure. Further experiments that examine changes in the binding interaction of FabI with substrate or acyl carrier protein after treatment with triclosan could reveal new insights into these conformational questions.

The selectivity exhibited by FabI for NADH over NAD^+^ can be plausibly explained based on the conformation of the nicotinamide-ribose glycosidic linkage observed in crystal structures (PDB codes 1QSG, 1DFI). In these and other structures containing NAD^+^, the C-O bond of the ribose-nicotinamide glycosidic linkage is rotated roughly perpendicular to the plane of the nicotinamide ring, which has been shown to be an unfavorable conformation for NAD^+^, but favorable for NADH.^28^ As a result, the affinity of the enzyme for NADH is expected to be much greater than its affinity for NAD^+^, which is confirmed by our experiments. However, this difference in affinity arising from the bound conformation of the coenzyme makes it difficult to rationalize a mechanism that explains the remarkable change in FabI affinity for NAD^+^ in the presence of triclosan. It appears to us that there are three possible explanations for these observations; we will present each explanation and the evidence in favor and against it.

First, it is possible that triclosan binding to FabI induces a conformational change that rotates the C-O glyosidic bond of NAD^+^ into a conformation that is parallel to the nicotinamide ring, thus creating a more favorable binding pose for NAD^+^. This does not seem very plausible since crystal structures containing both triclosan and NAD^+^ bound to the enzyme do not show any such change in the coenzyme conformation. For this explanation to be valid, we must assume that the crystal structures fail to accurately represent the solution-phase structure, which is unfounded in the absence of any corroborating evidence. A second explanation is that upon binding to FabI, triclosan directly interacts with both the enzyme and NAD^+^ in a way that creates a highly stable ternary complex, thus increasing the apparent affinity for NAD^+^. This is the traditional explanation and it is supported by apparent hydrogen bonding interactions between triclosan, NAD^+^ and FabI that are observed in the crystal structures. However, our experiments reported here cast some doubt on the plausibility of this explanation since triclosan completely dissociated from the enzyme even under gentle ionization conditions, while the NAD^+^ remains bound to the enzyme even under relatively high energy conditions (Figure 5). Although it is possible that our observations are merely an artifact of observing the enzyme in the gas-phase, our results remain difficult to harmonize with this traditional interpretation since no interactions that would be expected to stabilize gas-phase structures (e.g. salt bridges between the enzyme and ligands) are observed in the crystal structures. Finally, a third explanation is that triclosan binding to FabI induces conformational changes that are not observed in the crystal structure that result in highly stable interactions between the enzyme and NAD^+^. This explanation would provide the best fit with our observations in this study (both the CID fragmentation observations and microscopic K_A_ values), but would need more experimental evidence before it could be accepted as more plausible than the traditional hypothesis.

In the early 2000’s a series of papers from a team of scientists at GlaxoSmithKline reported a wide variety of FabI inhibitors, but despite their extensive efforts they were unable to discover any compounds with significantly greater inhibitory activity than triclosan.^29–32^ This failure appears remarkable in light of the extensive body of literature describing the details of FabI’s structure and function. In light of the results described in our study, it appears that this extensively studied enzyme-inhibitor system may still have secrets to share. It seems plausible that rigorously characterizing the conformational dynamics of FabI could reveal new opportunities for the design of future antibacterial therapies.

## Experimental Methods

### Instrumentation and reagents

MS analysis was performed on a Waters Synapt G2 Si QTOF mass spectrometer with a nanoESI inlet. NAD^+^, NADH and triclosan were purchased from Millipore-Sigma. A stock solution of 7 M ammonium acetate buffer was purchased from Thermo-Fisher and diluted to 500 mM for MS experiments.

### Protein preparation

Expression plasmids containing the *E. coli* FabI gene with an N-terminal His-tag were graciously provided by Dr. Charles O. Rock. Plasmids were transformed into BL21(DE3)-Gold *E. coli*, and six 1L bacterial cultures were grown overnight and then induced with IPTG for six hours. The cells in each culture were pelleted, frozen, and stored at - 20 C. Protein extracts were generated by thawing the frozen cell pellets and suspending the cells in 100 mL lysis buffer (50 mM TRIS pH 8, 300 mM sodium chloride, 10 mM imidazole, 1 mM PMSF, 1 mM EDTA, benzonase). The cell suspensions were sonicated on ice at 70% amplitude for 10 minutes (bursts of 10 seconds on, 10 seconds off). Cell debris was pelleted by centrifugation at 30,000 rcf for 25 minutes, and the supernatants from each extract were pooled and introduced to a Bio-Rad DuoFlow chromatography system with a 2 mL NiNTA column. The crude protein extract was fractionated using a 10-300 mM imidazole gradient in 50 mM TRIS, 300 mM sodium chloride buffer over 90 minutes with a flow rate of 1 mL/min. After initial purification by Ni-affinity chromatography, the collected FabI fractions were further purified by size-exclusion chromatography using a 30 mM PIPES (pH 8), 150 mM sodium chloride, 1 mM EDTA buffer. The final yield of FabI after expression and purification was 7 mg/mL. After purification, FabI was digested with thrombin to remove the His-tag and then directly analyzed by native MS. Protein concentration was determined from measurements of absorbance at 340 nM using a NanoDrop spectrophotometer, based on a monomeric molecular weight of 28,145 Da and a molar extinction coefficient of 16,180 M^−1^ cm^−1^.

### QTOF MS analysis

Samples were prepared at room temperature in 500 mM ammonium acetate and introduced into the MS using nanoESI borosilicate emitters (Thermo Fisher). A spray voltage of 1 kV was used with inlet backing pressure set to 0.2 bar and a cone voltage of 20 V and a source offset voltage of 40 V. The CID trap energy was set to 30 V and the trap helium flow was 4 mL/min.

The monomeric concentration of FabI was 5 µM in all experiments, based on absorbance measurements. For NADH binding affinity experiments, the concentration of NADH was varied in six samples at 10, 20, 40, 80, 160 and 320 µM. For NADH binding affinity experiments, the concentration of NADH was varied in six samples at 2, 4, 8, 16, 32 and 64 µM. Stock solutions of FabI, NAD^+^ and NADH were prepared in 500 mM ammonium acetate and diluted to the desired concentration. A 100 mM stock solution of triclosan was dissolved in DMSO and diluted to 1 mM in 1:4 DMSO:500 mM ammonium acetate to maintain solubility of the ligand. The triclosan was diluted to the final target concentrations in mixtures prepared in 500 mM ammonium acetate, leaving the final concentration of DMSO <0.5%.

### Spectral processing and visulization

For spectra shown in figures, each spectrum was processed using a background subtraction function and a smoothing function in Waters MassLynx (ver 4) software.

### Data analysis and calculation of binding affinities

We created a series of R-scripts to extract peak intensity values from batches of individual spectra (scripts included in Supporting Information). Briefly, the RAW files from the Synapt instrument were first converted to the mzML format using ProteoWizard^33^ and then imported into the MALDI-QUANT package^34^ in R^35^ using RStudio^36^. MALDI-QUANT was used to smooth and average the spectra from each experiment, and then peak intensities were identified and written to a feature matrix. The feature matrix was then imported into a spreadsheet to calculate binding affinities.

Binding affinities were calculated using the formulas described by J. Klassen and colleagues.^22^ Specifically, the macroscopic *K*_*A*_ for each experiment was calculated using the formula

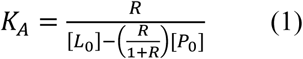

where *R* is the ratio of the sum of the intensities of all peaks in the mass spectrum corresponding to ligand-bound protein to the intensities of the peaks corresponding to free protein, *L*_*0*_ is the initial concentration of the ligand and *P*_*0*_ is the initial concentration of protein. Values from all charge states were combined into each value of *R* to account for all of the protein represented in each spectrum. This formula was used instead of fitting the data to a non-linear formula because it was consistent with the formula used to calculate the microscopic affinities.

To calculate the microscopic affinities, the formula

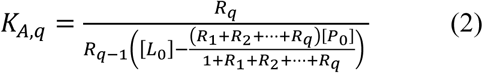

was used, based on the expected expression for the sequential binding each ligand

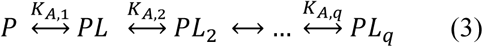

In equation 2, *R*_*q*_ corresponds to the ratio of the intensity of the peak representing *q* ligands bound compared to the free protein and *L*_*0*_ and *P*_*0*_ are the same as described for equation 1.

These equations generate a macroscopic *K*_*A*_ value and up to four microscopic *K*_*A*_ values for each experiment where ligand concentration was varied. The macro- and microscopic *K*_*A*_ values for each titration series were averaged (mean) to account for differences arising from changing ligand concentrations. Each titration series was performed in triplicate, and the mean macro- and microscopic *K*_*A*_ values were used to calculate the values shown in Table 1. The statistical relationship that is expected to exist between equivalent, independent ligand binding sites is given by Klassen and colleagues as

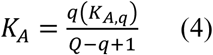

where Q is the number of ligand binding sites.

## Acknowledgements

Support from the US National Institutes of Health (R01GM103479, S10OD018504 to J.A.L.) and the US Department of Energy (DE-FC02-02ER63421) are gratefully acknowledged.

