## Supporting Information for "Characterization of protein-ligand binding interactions of enoyl-ACP reductase (FabI) by native MS reveals allosteric effects of coenzymes and the inhibitor triclosan"

#### Supplementary Figures

**Figure S1.** Denatured spectrum of FabI

**Figure S2.** MSMS spectra of the FabI apo enzyme

**Figure S3.** MSMS spectra of the FabI treated with $NAD^+$ and triclosan

**R script for extracting peak intensities from spectra**

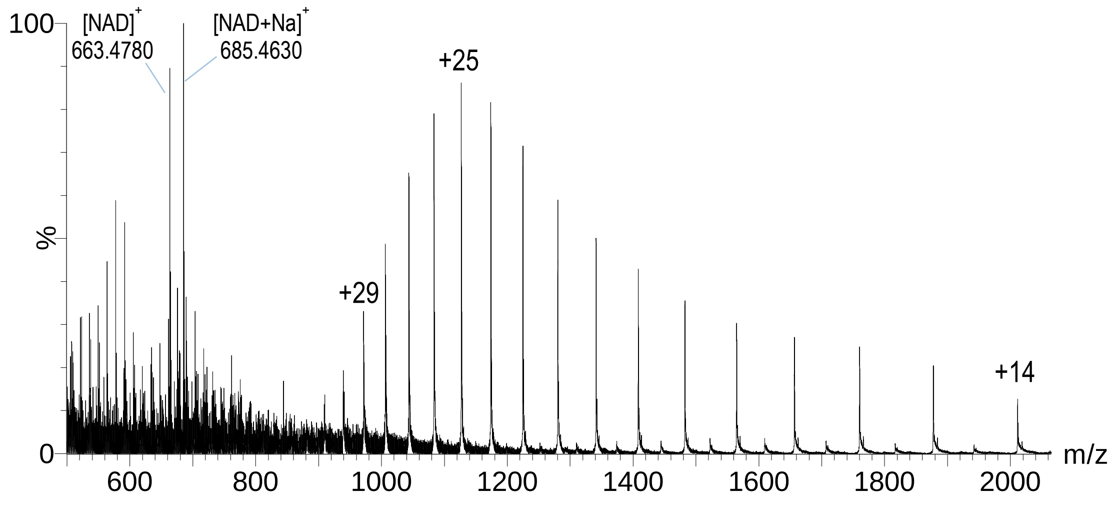


**Figure S1.** Denatured spectrum of FabI. Charge states are shown for the protein peaks; peaks corresponding to $NAD^+$ and $[NAD+Na]^+$ are labeled. The observation of the $NAD^+$ peaks confirms the identity of the small FabI-ligand complex observed in the native MS spectrum of purified FabI.

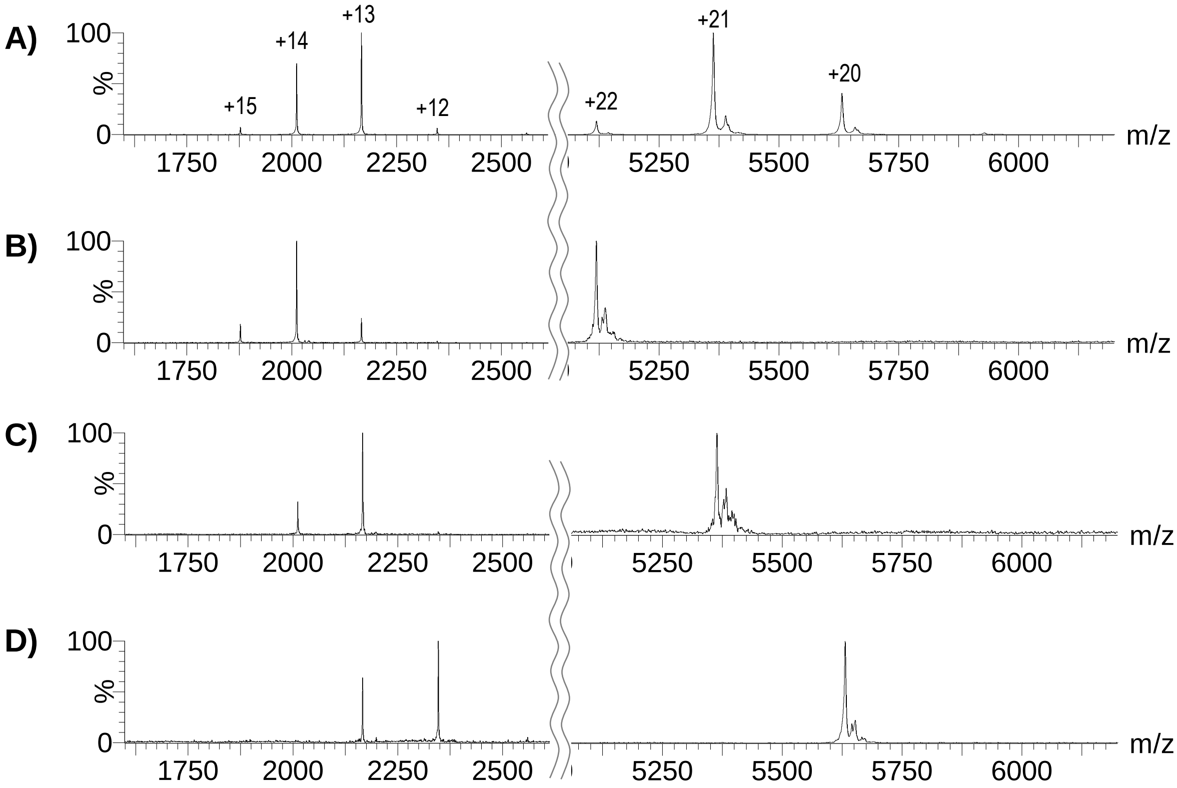


**Figure S2.** MSMS spectra of the FabI apo enzyme (5 µM). A) CID 90V spectrum showing the tetramers and monomeric dissociation products, with charge state shown above each peak. B) MSMS fragmentation spectrum of the m/z 5119 peak. The +22 tetramer primarily yields the +14 monomer. C) MSMS fragmentation spectrum of the m/z 5363 peak. The +21 tetramer primarily yields the +13 monomer. B) MSMS fragmentation spectrum of the m/z 5631 peak. The +20 tetramer primarily yields the +12 monomer.

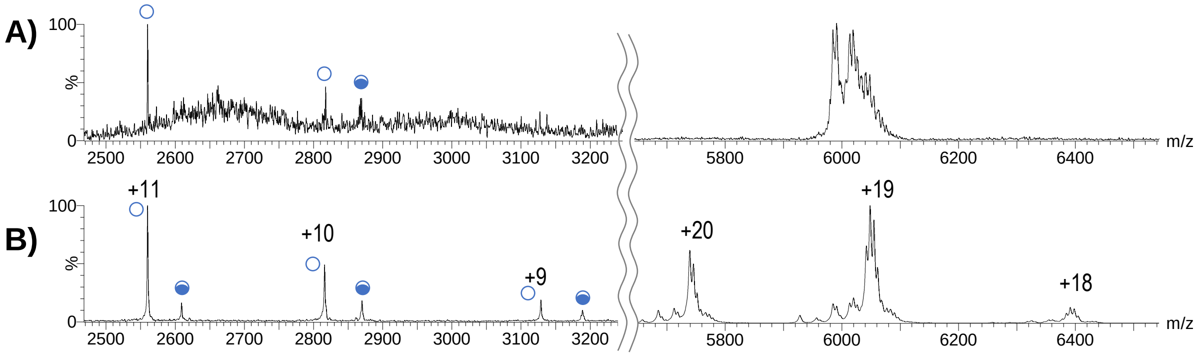


**Figure S3.** MSMS spectra of the FabI (5 µM) treated with $NAD^+$ (30 µM) and triclosan (16 µM). A) MSMS fragmentation spectrum of the m/z 6048 peak. The +19 FabI-$NAD^+_4$ complex yields monomeric dissociation products in the +11 and +10 charge state with a FabI-ligand monomeric complex observed in the +10 charge state. B) CID 90V spectrum showing the tetramers and monomeric dissociation products, with charge state shown above each peak. Open circles designate monomers with no ligand; partially filled circles designate monomers with a bound $NAD^+$ fragment.

### R script for extracting peak intensities from spectra

```
# Using MALDIquant and nls to calculate Ka/Kd values for NADH titration
# Created by Matt Joyner July 6, 2019

# load MALDIquant libary
library("MALDIquant")
library("MALDIquantForeign")

# set directory to folder containing files for conversion
setwd("INSERT_PATH_TO_DATA")

# import avgeraged spectrum of each point in triplicat concentration curves
#from  June 11, 2019 (p. 189)
#replicate 1
spec189a <- import("avg_189a.mzML")
spec189b <- import("avg_189b.mzML")
spec189c <- import("avg_189c.mzML")
spec189d <- import("avg_189d.mzML")
spec189e <- import("avg_189e.mzML")
spec189f <- import("avg_189f.mzML")

#from June 11, 2019 (p. 189)
#replicate 2
spec189j <- import("avg_189j.mzML")
spec189k <- import("avg_189k.mzML")
spec189l <- import("avg_189l.mzML")
spec189m <- import("avg_189m.mzML")
spec189n <- import("avg_189n.mzML")
spec189o <- import("avg_189o.mzML")

#from June 13, 2019 (p. 191)
#replicate 3
spec191a <- import("avg_191a.mzML")
spec191b <- import("avg_191b.mzML")
spec191c <- import("avg_191c.mzML")
spec191d <- import("avg_191d.mzML")
spec191e <- import("avg_191e.mzML")
spec191f <- import("avg_191f.mzML")

# combine individual spectra into single list
nadhspec <- list(spec189a[[1]], spec189b[[1]], spec189c[[1]], spec189d[[1]],
spec189e[[1]], spec189f[[1]], spec189j[[1]], spec189k[[1]], spec189l[[1]],
spec189m[[1]], spec189n[[1]], spec189o[[1]], spec191a[[1]], spec191b[[1]],
spec191c[[1]], spec191d[[1]], spec191e[[1]], spec191f[[1]])
names(nadspec) <- c("spec189a", "spec189b", "spec189c", "spec189d",
"spec189e", "spec189f", "spec189j", "spec189k", "spec189l", "spec189m",
"spec189n", "spec189o", "spec191a", "spec191b", "spec191c", "spec191d",
"spec191e", "spec191f")
nadhspec

# create plot of spectra to visually confirm quality
plot(nadhspec[[1]], ylim = c(0,1500), xlim = c(5300, 5600))
```

```
lines(nadhspec[[7]], col = c("blue"))
lines(nadhspec[[13]], col = c("red"))
lines(nadhspec[[4]], col = c("green"))

# create feature matrix from spectra by smoothing spectra, calculating
single:noise, finding peaks, binning spectra and then creating feature matrix
nadhsmooth <- smoothIntensity(nadhspec, method = "SavitzkyGolay",
halfWindowSize = 10)
noise <- estimateNoise(nadhsmooth[[1]])
lines(noise[,1], noise[,2]*.3, col = "orange")
nadhpeaks <- detectPeaks(nadhsmooth, method = "MAD", halfWindowSize = 200,
SNR = 0.3)
nadhbins <- binPeaks(nadhpeaks, tolerance = 0.002)
nadhfilt <- filterPeaks(nadhbins, minFrequency = 0.1) # minFrequency set to
remove peaks that only appear in less than 2 spectra
nadhfeatmatrix <- intensityMatrix(nadhfilt, nadhsmooth)
head(nadhfeatmatrix[,3:8])

#rename columns to easier to use names (e.g. truncated m/z values)
peaknames <-
substr(colnames(nadhfeatmatrix)[1:length(colnames(nadhfeatmatrix))], 1, 4)
peaknames
colnames(nadhfeatmatrix) <- peaknames
head(nadhfeatmatrix[0,3:8])
write.csv(nadhfeatmatrix, file = "nadhplusfeaturematrix.csv")
```
